# A sweep of earth’s virome reveals host-guided viral protein structural mimicry; with implications for human disease

**DOI:** 10.1101/2020.06.18.159467

**Authors:** Gorka Lasso, Barry Honig, Sagi D. Shapira

## Abstract

Viruses deploy an array of genetically encoded strategies to coopt host machinery and support viral replicative cycles. Molecular mimicry, manifested by structural similarity between viral and endogenous host proteins, allow viruses to harness or disrupt cellular functions including nucleic acid metabolism and modulation of immune responses. Here, we use protein structure similarity to scan for virally encoded structure mimics across thousands of catalogued viruses and hosts spanning broad ecological niches and taxonomic range, including bacteria, plants and fungi, invertebrates and vertebrates. Our survey identified over 6,000,000 instances of structural mimicry, the vast majority of which (>70%) cannot be discerned through protein sequence. The results point to molecular mimicry as a pervasive strategy employed by viruses and indicate that the protein structure space used by a given virus is dictated by the host proteome. Interrogation of proteins mimicked by human-infecting viruses points to broad diversification of cellular pathways targeted via structural mimicry, identifies biological processes that may underly autoimmune disorders, and reveals virally encoded mimics that may be leveraged to engineer synthetic metabolic circuits or may serve as targets for therapeutics. Moreover, the manner and degree to which viruses exploit molecular mimicry varies by genome size and nucleic acid type, with ssRNA viruses circumventing limitations of their small genomes by mimicking human proteins to a greater extent than their large dsDNA counterparts. Finally, we identified over 140 cellular proteins that are mimicked by CoV, providing clues about cellular processes driving the pathogenesis of the ongoing COVID-19 pandemic.

## INTRODUCTION

Viruses deploy an array of genetically encoded strategies to coopt host machinery and support viral replicative cycles. Among the strategies, protein-protein interactions, mediated by promiscuous, multifunctional viral proteins are widely documented. Targeted discovery tools, focused largely on viruses of public-health importance, have experimentally mapped thousands of virus-host protein complexes and structure-informed prediction algorithms have allowed discovery of such interactions across all fully sequenced human infecting viruses^1^. Molecular mimicry, manifested by structural similarity between viral and endogenous host proteins, allow viruses to harness or disrupt cellular functions including nucleic acid metabolism and modulation of immune responses. Yet, while examples of this latter strategy pepper the literature^2-4^, most have focused on human infecting viruses^5,6^ and a systematic analysis of pathogen-encoded molecular mimics has not been performed.

The vast genomic landscape occupied by viruses hampers the discovery of evolutionary relationships between viral proteins and their hosts. As is well known however, since 3-dimensional (3D) protein structure is much better conserved than sequence, structural information can be used to interrogate evolutionary relationships^1,7^ as well as uncover virus-encoded structural mimics that cannot be detected by sequence relationships (see Methods). Here, we use protein structure similarity to identify virally encoded mimics of host proteomes. Briefly, we first employ sequence-based methods to identify proteins that have similar structures to queried viral proteins and then use structural alignment to find “structural neighbors” of viral proteins (Figure 1a). We refer to the corresponding viral proteins as mimics of their host-encoded neighbors. We applied the approach to a set of 337,493 viral proteins representing 7,486 viruses across a broad host taxonomic range, including bacteria, plants and fungi, invertebrates and vertebrates. Our survey identified over 6,000,000 instances of structural mimicry, the vast majority of which (>70%) cannot be discerned through protein sequence alone (see below). Our results point to molecular mimicry as a pervasive strategy employed by viruses and indicate that the protein structure space used by a given virus is dictated, at least in part, by the host proteome.

**Figure 1.**
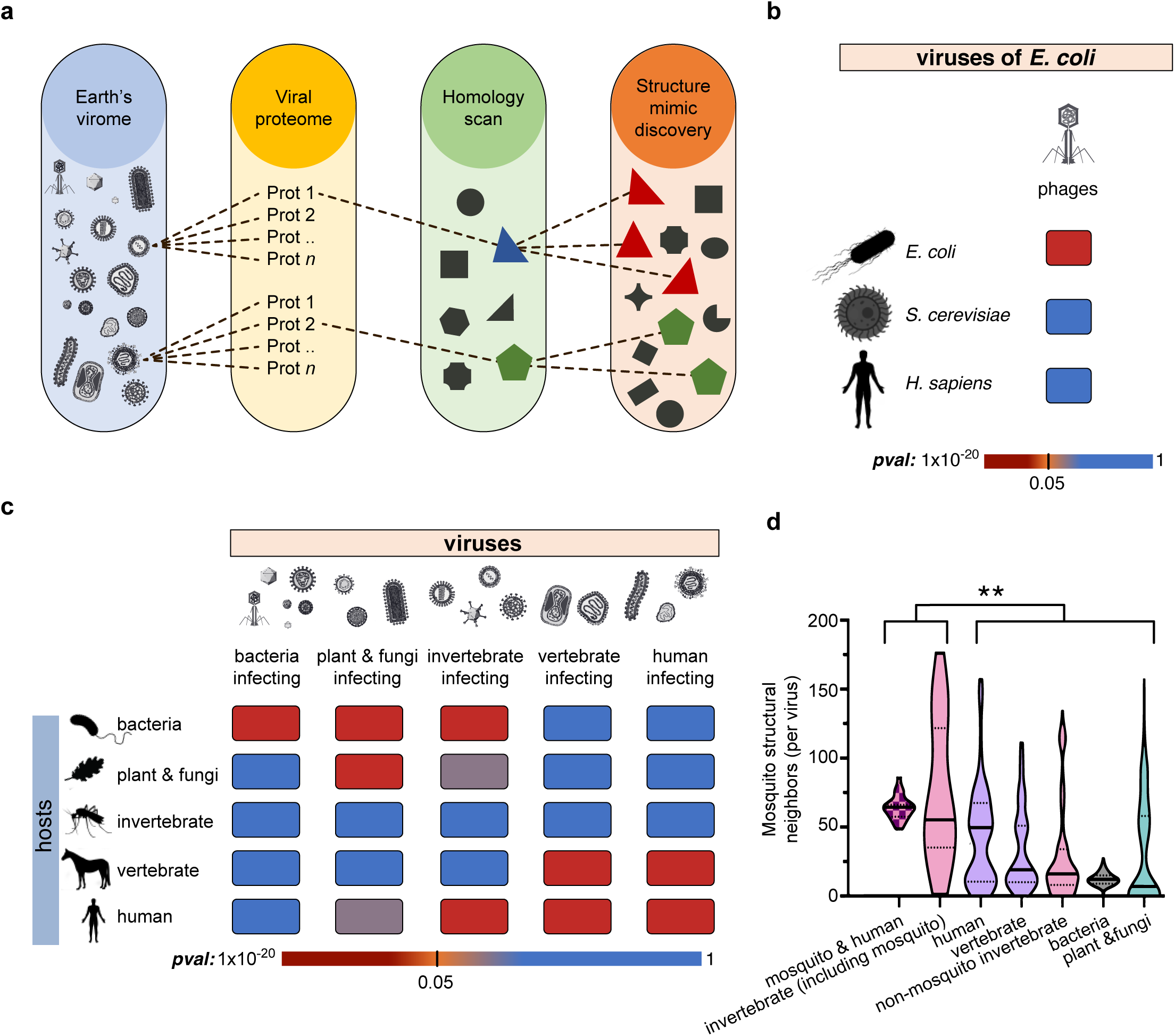
Identification and characterization of virus-host structural relationships reveals utilization of structure space guided by host taxonomic division. **a**, Graphical schematic of the strategy implemented to reveal virus encoded protein mimics. *Sequence-based* searches are first used to identify proteins in the PDB that are structurally related to viral proteins. These intermediate, sequence-detected structural templates are then used, through *structural alignment*, to search the PDB for proteins that are geometrically similar to viral proteins – thus revealing viral mimicry **b**, Significance of overlap, measured in terms of numbers observed relationships (where SAS < 2.5Å) between the structure space of *E*. coli infecting phages and *E. coli, S. cerevisiae*, and human proteomes (See Methods for details). **c**, Significance of overlap, measured in terms of numbers of observed relationships (where SAS < 2.5Å) between the structure space of all catalogued viruses and hosts in different taxonomic divisions. **d**, Distribution of structural neighbors of the modeled mosquito proteome for different virus classes (brackets denote distributions used in hypergeometric test, see methods; \*\**P*value < 0.0001).

We further observe that the manner and degree to which viruses exploit molecular mimicry varies by genome size and nucleic acid type. For example, while human-infecting and arthropod-infecting viruses occupy a structure space most similar to their host proteome, arboviruses, which are transmitted to humans by insect vectors, encode promiscuous proteins that mimic both human and insect proteins. In addition, we find that, relative to their proteome size, ssRNA viruses, including coronaviruses (CoV), have circumvented the limitations of their small genomes by mimicking human proteins to a greater extent than their large dsDNA counterparts like Pox- and Herpes-viruses. Interrogation of proteins mimicked by human-infecting viruses points to broad diversification of cellular pathways targeted via structural mimicry, identifies biological processes that may underly autoimmune disorders, and reveals virally encoded mimics that may be leveraged to engineer synthetic metabolic circuits or may serve as targets for therapeutics. Finally, we identified over 140 cellular proteins (including members of the complement activation pathway and critical regulators of innate and adaptive immunity) that are mimicked by CoV, providing clues about the cellular processes underlying the pathogenesis driving the ongoing COVID-19 pandemic.

## Results

### Mining the virome-wide structure space to identify host proteins mimicked by viruses

In order to identify mimicry relationships, we implemented the strategy summarized in Figure 1a. Sequence-based searches (see Methods) of 337,493 viral proteins (269,077 unique sequences) against the PDB identified structural templates for 92,868 (34.5%), with large disparities in coverage: structural templates were identified for 64.7% of the proteins from human-infecting viruses while the coverage for non-vertebrate-infecting viruses ranged from 31% to 38% (Figure S1a; Table S2). As shown in Figure S1b, structural templates for vertebrate and human infecting viruses came primarily from other viruses while, for example, bacterial proteins provided most of the templates for bacteria-infecting viruses. Most of the templates for invertebrate-infecting viruses came from vertebrate and bacterial proteins, likely reflecting the limited structural coverage of non-vertebrate viruses and invertebrate hosts in the PDB. Ska was used To measure structure similarity between protein structures, we utilized Ska, an extensively utilized and validated tool for inference of structure-based functional relationships even in the absence of detectable sequence similarity^1,8-13^. In addition to Ska, we employed a conservative global structural similarity criteria (Structural Alignment Score^14-16^, SAS < 2.5Å) to infer structural mimics and minimize biases imposed by local structural similarities (see Methods). As shown in Figure S2, this approach enables rediscovery of known virus encoded protein structure mimics. Structural alignment of the 92,868 templates with PDB proteins identified 6,083,167 mimicry relationships, involving 88,715 viral proteins and 26,542 host protein structures present in the PDB (data is available for download at https://github.com/gorkaLasso/VirusHost_mimicry). The vast majority of these mimicry relationships (72.2%) could not be retrieved through sequence similarity alone based on the requirement that no e-value obtained from any one or three sequence-based methods was greater than 1×10^−6^ (see Methods).

### Host range and taxonomic division drives viral protein structure space and mimicry

Taxonomic enrichment analysis (Figure 1b-c) reveals a clear correlation between the extent of virus structural mimicry and the identity of the host. Since their proteomes are well represented in the PDB, we first focused on the proteomes of *E. coli*-infecting phages, their natural host *E. coli*, the budding yeast *Saccharomyces cerevisiae*, and human. Significant structure similarity is only observed between *E. coli*-infecting phages and their natural host (Figure 1b). As shown in Figure 1c, with the exception of invertebrate infecting viruses which constitute a special case (see below), a broader analysis across all 7,486 catalogued viruses and 38,363 proteins from 4,045 putative hosts demonstrates that every virus group exhibits significant protein structural similarity to the taxonomic division of their hosts. In addition, viruses exhibit structural similarity to proteomes in other taxonomic divisions that are close in evolutionary distance to the host taxonomic division, and reduced structural similarity to more distant taxonomic divisions (e.g. the structural space of human-infecting viruses are enriched for human, vertebrates and invertebrate proteins but not bacteria or plant proteins, Figure 1c). Such similarities are reflected in evolutionary relationships between different taxonomic divisions. For instance, accumulating evidence describes some compartments of the phytosphere as environmental host spots for horizontal gene transfer (HGT) that facilitate genetic exchange between plants and bacteria^17^, underscoring the significant structural relationship between plant and fungi-infecting viruses and phages (Figure 1c). Moreover, in agreement with the results obtained for the broader virus categories, we found the protein structure space of human-infecting viruses to be enriched with structures found in human and non-human vertebrate proteomes (Figure 1c) – pointing to a possible path through which zoonotic infections may jump from animal reservoirs to cause human disease.

Unlike viruses of bacteria and vertebrates, we do not observe significant structural similarity between proteomes of invertebrate-infecting viruses and their invertebrate hosts – perhaps owing to the fact that invertebrates account for the taxonomic group with the lowest structural coverage in the PDB (containing just 2,687 unique invertebrate proteins). To bridge this knowledge gap, we modeled the proteome of the mosquito *Ae. Aegypti* (see Methods) and compared the protein structure space of mosquito with each group of viruses. As shown in Figure 1d, we find that mosquito-infecting viruses occupy a significantly larger mosquito structural space than viruses that infect hosts in other taxonomic divisions. Moreover, we observe that mosquito-borne arboviruses, human-infecting viruses that use mosquitoes as vectors, engage a larger space of mosquito protein mimics than human-infecting viruses that do not use mosquitoes as vectors (*P*value 2.3×10^−3^; Figure 1d). Together, these data further highlight the evolutionary pressures imposed on viruses to utilize a protein structure space defined by their host’s range.

### Diversification of mimicry strategies among human-infecting viruses

Figure 2a displays the number of human structural mimics for different groups of viruses. A number of viral families, including *Hepadnaviridae, Bunyaviridae, Papillomaviridae, Parvoviridae, Retroviridae, Arenaviridae* and *Polyomaviridae*, utilize structural mimicry far less than others. On the other hand, poxviruses and herpesviruses, large dsDNA viruses that are less constrained by genome size (Figure 2b) and more likely to retain horizontally transferred genes^3,5^, encode many human protein structure mimics (Figure 2a). In contrast, relative to their proteome size, small RNA viruses, which are constrained by genome size, display enhanced structural promiscuity, with a larger number of human protein mimics per residue (Figure 2c). So, while viruses constrained by genome size may be restricted in their ability to expand their genomes through horizontal gene transfer, they have circumvented this limitation by encoding multifunctional proteins that mimic domains across multiple host proteins. These observations underscore the role of structural mimicry in evolving expanded molecular functionality in size constrained genomes.

**Figure 2.**
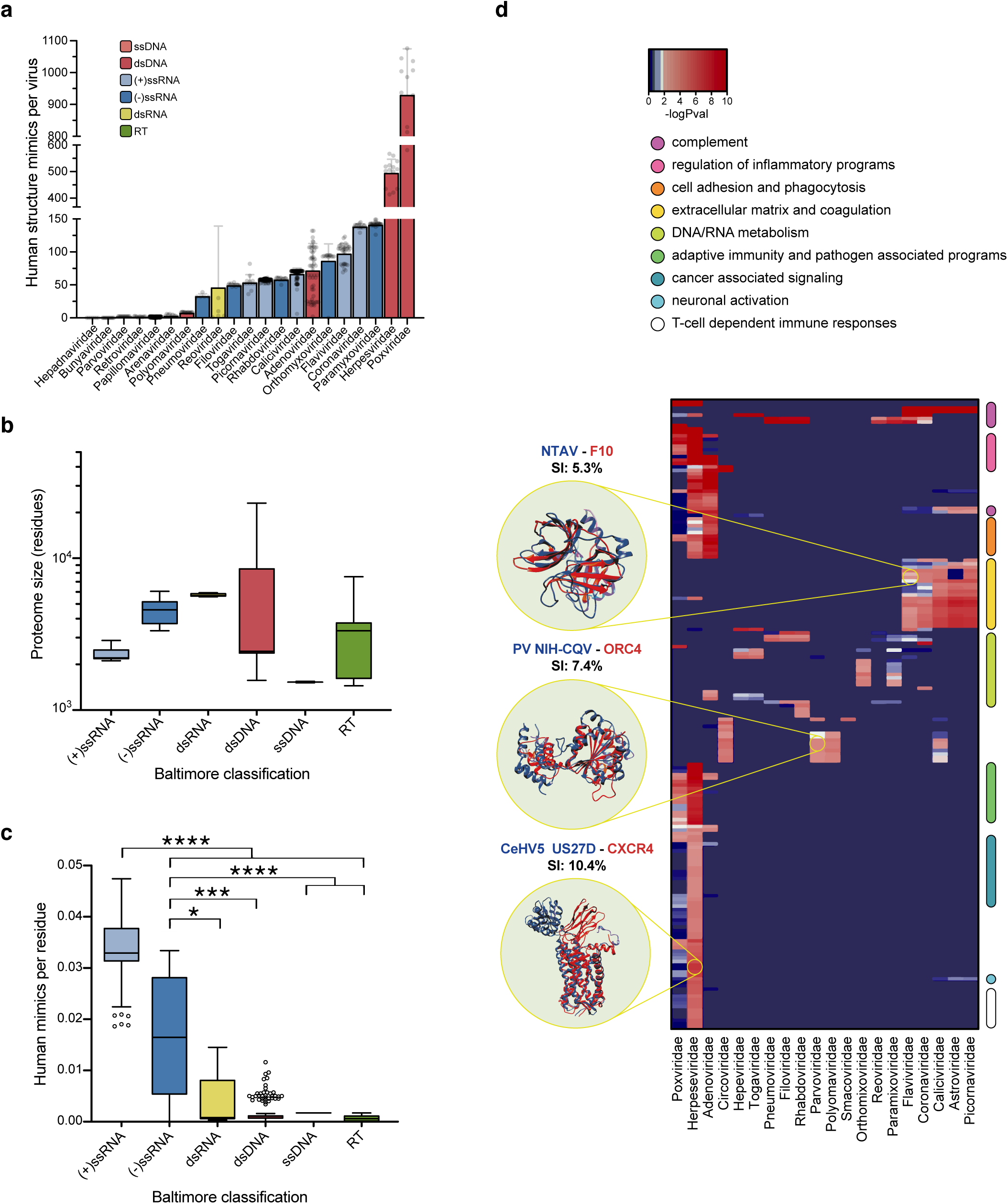
Mimicry strategies utilized by human-infecting viruses vary by nucleic acid type and proteome size. **a**, Number of virus-encoded human-protein structure-mimics for human-infecting viruses, grouped by viral family. **b**, Virus proteome sizes (amino acids) grouped by nucleic acid type. **c**, Number of structural mimics encoded per residue for human-infecting viruses, grouped by nucleic acid type. Virus family classification and nucleic acid type are as indicated by color key and labels (**a**-**c;** box and whisker plots **b** and **c** indicate lower and upper quartiles and span minimum and maximum values). **d**, Hierarchical clustering of enriched biological pathways (rows) targeted through mimicry by human infecting virus families (columns). Biological pathway categories are highlighted by color bars on the right of the heat-map. Examples of structural alignments between virus encoded protein mimic (red) and human protein (blue); sequence identities (SI) are indicated in black; *P*values indicated by color key.

### Pathways targeted by viral mimicry

#### Human-infecting DNA encoded viruses

The critical nature of virus-host PPIs in engaging host molecular machinery to fulfill viral life cycles has led to convergence of evolutionarily unrelated viruses around key cellular pathways (e.g. human-infecting DNA, RNA and retro-transcribing viruses converge, via different sets of PPIs, on multiple cellular metabolic pathways)^1^. In contrast, as shown in Figure 2d and Table S3, structurally mimicked proteins revealed little such convergence. Moreover, when convergence was observed, it often involved 2 or 3 viral families belonging to the same Baltimore classification. For example, owing to their large genomes and in agreement with their propensity to acquire host genes through HGT, *Herpesviridae, Adenoviridae* and *Poxviridae*, families of dsDNA viruses are enriched for the largest number of mimicked biological pathways, 113, 38 and 35 respectively (Figure 2d and Table S3). Yet, while these viruses harbor a large number of human protein mimics, they do not mimic cellular components of RNA and DNA metabolism – likely reflecting the fact that they encode polymerases (as is the case of *Poxviridae*) and/or integrate into the cellular genome and are therefore bystander participants in cellular DNA replication.

We observe an enrichment of cytokine related pathways in poxvirus- and herpesvirus-mimicked proteins, and chemokine-related pathways in proteins mimicked by herpesviruses (Table S3 and Figure 2d). Indeed, poxviruses and herpesviruses have evolved a repertoire of immune evasion strategies by acquiring immunomodulatory host genes through genetic recombination (often involving cytokines and cytokine receptors). Undetectable sequence similarity with the host has complicated past attempts to identification virus-host relationships^6,18^. However, while many of the relationships between herpesvirus and chemokine or chemokine receptors can be discerned from sequence, we have expanded the list of known mimics to include 61 new herpesviral structure mimics not discernible from sequence alone (Figure 2d; complete list of mimics is available at https://github.com/gorkaLasso/VirusHost_mimicry). For example, as shown in Figure 2d, we observed that US27D protein from Cercopithecine betaherpesvirus 5, a cytomegalovirus, mimics human CXCR4 (sequence identity: 10.4%), a chemokine receptor that regulates T-cell inflammatory programs and can serve as a co-receptor for HIV^19^. It is unclear if the structural mimics encoded by DNA viruses that we have identified are a result of sequence divergence of products acquired by HGT or through convergent evolution. Yet, they highlight the pervasive use of mimicry by DNA viruses and point to novel host factors that are targeted by structural mimicry.

#### Human-infecting RNA encoded viruses

As highlighted above, RNA viruses have circumvented limitations imposed on their small proteomes by encoding multifunctional and structurally promiscuous proteins that mimic domains across multiple host proteins (Figure 2c). In accordance with known phenotypes (like fibrinolysis and hemorrhagic fever) associated with Dengue virus (DENV; *Flaviridae*) infection^20,21^, we find that proteins mimicked by these RNA viruses are enriched for functions and pathways related to blood coagulation and the complement pathway – an observation that is mirrored by PPIs mediated by proteins encoded by these viruses (we previously found that PPI networks mediated by proteins of 51 of 56 flaviviruses were enriched for “complement and coagulation cascades”)^1^. While examples of such mimicry have been identified through sequence homology between DENV C, prM, E and NS1 proteins and human coagulation factors^20,21^, we find similar mimicry, that cannot be discerned through sequence, employed by four other (+)ssRNA virus families (168, 166, 1376 and 1293 structural relationships between coagulation factors and astroviridae, coronaviruses, caliciviruses and picornaviruses, respectively; sequence identity < 20%, Table S3, data available at https://github.com/gorkaLasso/VirusHost_mimicry). So, while DENV immunization or exposure elicits antibodies that cross-react with multiple coagulation factor-associated pathological states^20,21^, our observations suggest that infections with viruses belonging to these other families may also result in similar imbalances in immune responses. Indeed, viruses within these families (e.g. SARS-*Coronaviridae*, Hepatitis A-*Picornaviridae* and Hepatitis C-*Flaviridae*), are known to be associated with coagulation disorders such as thrombocytopenia, thrombosis and hemorrhage^22^.

### Coronaviruses

Knowledge of the precise regulatory programs that control the viral life cycle and mediate immune pathology can provide valuable clues about disease determinants. A closer look at coronaviruses underscores the level of diversification and structural promiscuity of protein mimics encoded by these viruses. As highlighted in Figure 2c, relative to proteome size, positive sense single stranded RNA ((+)ssRNA) viruses including CoV utilize structural mimicry more so than other human-infecting viruses. We identified 145 cellular proteins that are mimicked by three or more coronaviruses (data available at https://github.com/gorkaLasso/VirusHost_mimicry). Among these, we have identified CoV-encoded structural mimics that target key regulators of immune anti-viral programs.

First, in the context of CoV infection, intracellular sensing of viral RNA through DDX58 (also known as RIG-I; Retinoic Acid-Inducible Gene I), a pathogen recognition receptor, triggers a signaling cascade that results in the production and secretion of Type I interferons (IFNα/β) ^23,24^. IFNα/β in turn binds to cell surface receptors (IFNRs) on nearby cells and, following a signal-transduction cascade, leads to STAT1-mediated (<u>S</u>ignal <u>T</u>ransducer And <u>A</u>ctivator Of <u>T</u>ranscription 1) upregulation of hundreds of IFN-stimulated genes (ISGs) which govern cellular responses to infection and render cells refractory to viral infection. These responses are indispensable for control of viral replication and initiation of long-term immune responses ^23,24^. As shown in Figure 3, we find that NSP13 of CoV mimics DDX3X and RIG-I as well as other helicase proteins, critical cellular components that cooperate through both direct and indirect interactions to initiate immune responses to viral infection. Importantly, RIG-I ligands include ssRNAs that contain a 5’-triphosphate moiety^23,24^ and previous biochemical characterizations have attributed multiple enzymatic activities to NSP13, including hydrolysis of NTPs and dNTPs and RNA 5’-triphosphatase activity^25,26^. So, mimicry of RIG-I may serve to reduce the amount of viral RNA that is available to be sensed in infected cells. In addition, the CoV replicase complex has recently been demonstrated to interact with host cell DDX proteins^27,28^, suggesting that coronaviruses may utilize both PPIs and structural mimicry to manipulate functions of these proteins.

**Figure 3.**
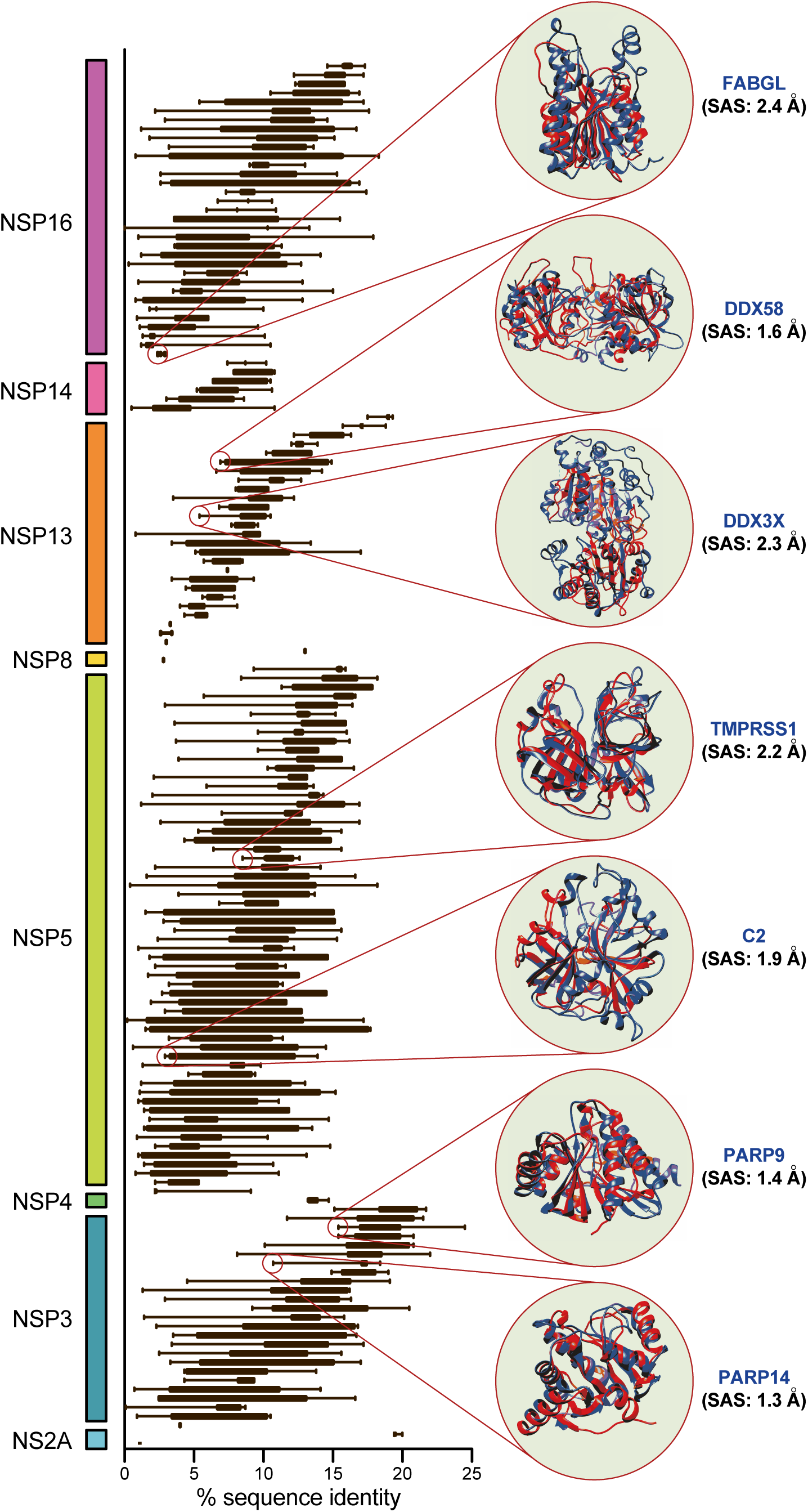
Coronavirus encoded proteins mimic known regulators of human immune response to infection. A total of 145 human proteins are structurally mimicked by 8 CoV proteins (grouped and indicated by color bars). Shown are box and whisker plots indicating lower and upper quartiles and spanning minimum and maximum pairwise % sequence identities between human and virus encoded proteins (variability in sequence identity reflects CoV specific sequence divergence; each human protein is mimicked by ≥3 viruses). Highlighted, are seven examples of virus encoded human protein mimics (virus protein structure in red; human protein structure and gene symbol in blue; SAS indicated in black).

In addition to mimicking critical components of the cellular nucleic acid sensing machinery, we have identified CoV-encoded structural mimics that target key regulators of downstream anti-viral programs mediated by STAT1. As shown in Figure 3, we observe significant structural similarity between CoV NSP3 protein and human PARP9 and PARP14, ADP-ribosyltransferases with known cross-regulatory roles in controlling STAT1-mediated signaling^29^. Though the precise mode of action has remained unresolved, CoV ORF3, which encodes NSP3, has indeed been shown to be an antagonist of IFNα/β signaling and interferes with STAT1nuclear translocation^30^. Our results point to structural mimicry as a molecular explanation underlying these observations.

Lastly, as discussed above, we find that structural mimicry of complement components is a feature shared across all coronaviruses that were part of this study (including SARS-CoV-2). The complement system is a critical regulator of immune responses to microbial threats, but when dysregulated by age-related effects or excessive acute and chronic tissue damage, complement activation can contribute to pathologies mediated by inflammation. Our data implicates virally encoded structural mimics that may contribute to SARS-(*Coronaviridae*) associated coagulation disorders like thrombocytopenia, thrombosis and hemorrhage^22^ – disease outcomes shared with Hepatitis A (*Picornaviridae*) and Hepatitis C (*Flaviridae*) infections (which also encode coagulation and complement cascade mimics). In addition, while the age-related differences in susceptibility to CoV (including SARS-CoV-2) are likely a consequence of multiple underlying variables, one possibility is that prior exposure to coronavirus may potentiate complement-mediated responses to subsequent coronavirus infections. Moreover, a corollary of these observations is that complement-associated syndromes (including complement deficiencies or hyper-activation phenotypes like those associated with macular degeneration) may impact clinical outcome of SARS-CoV-2 infection. Indeed, in the as part of a separate retrospective clinical study, we have demonstrated that history of macular degeneration, is a significant risk factor for morbidity and mortality in SARS-CoV-2 infected patients – effects that could not be explained by age or sex^31^.

## DISCUSSION

The mimicry of host protein structures is a strategy used by viruses to harness or disrupt host cellular functions^2-6^. Structural mimicry can occur at the level of entire protein domains or in the form of “interface mimicry”, where the structure of host protein residues involved in protein-protein interactions is mimicked on the surface of a viral protein^32-34^. High throughput experimental methods, focused largely on viruses of public-health importance, have experimentally mapped thousands of virus-host protein complexes while structure-based computational tools based on interface mimicry have reported notable successes in the prediction of protein-protein interactions, PPIs^32-34^. Recently we reported a structure-informed prediction algorithm, P-HIPSTer, that exploits domain level similarity and interface properties to predict virus/host PPIs across all fully sequenced human infecting viruses^1^. Studies such as these have revealed valuable information about the molecular mechanisms that underlie viral infection. Yet, while most of these experimental and computational studies have focused on PPIs, a systematic analysis of viral mimicry of host proteins has not been reported. Here, rather than considering PPIs, we explore the extent to which viral mimicry of host proteins is a common phenomenon and ask whether the “structure space” occupied by a particular virus is related to that of the host it infects. Moreover, by analyzing the host proteins that are mimicked we are able to gain functional insights about infection mechanisms that are complementary to those gained by predicting PPIs.

Structure enables identification of mimics between viral and host proteins that cannot be observed from sequence alone. Mimicry relationships have been detected through sequence similarity and linear motif co-occurrence. We bypass limitations of sequence-based approaches by leveraging 3D protein structure to identify virally encoded proteins with significant structural similarity to host proteins. Although examples of pathogen-encoded protein mimicry have been reported in the literature, to our knowledge, this is the first comprehensive and systematic analysis of molecular mimicry across the earth’s virome – spanning 337,493 viral proteins from 7,486 viruses (bacteria, plant-fungi, invertebrate and vertebrate-infecting viruses) and 38,363 proteins from 4,045 putative hosts.

Our pipeline uses a conservative global structural similarity criteria (Structural Alignment Score, SAS < 2.5Å) to infer structural mimics^14-16^. Consequently, our approach does not consider local structural similarities defined by small protein regions and linear motifs (frequently found in disordered regions)^4,32^. Importantly, structural mimicry does not necessarily imply functional mimicry. Furthermore, some of the mimics that we identify may represent spurious structural relationships that stem from underlying functional class of a give pair of proteins (virus and host encoded proteases). Future experimental and functional interrogation will be necessary to demonstrate functional implications for the molecular mimics discovered as part of this work. Nevertheless, the results illustrate the ability of protein structure-based analysis to infer structural, functional and evolutionary relationships between viruses and their hosts.

Our results demonstrate that regardless of genome size, replicative cycle, or ecological niche, and the evolutionary pressures imposed on viruses is reflected in the structure space they occupy and illustrate that mimicry may both constrain and enable host range. Of note, while structural similarity between viral and host proteins serves as a scaffold enabling viral proteins to coopt host pathways, amino acid identities at key functional sites will ultimately shape the extent to which host pathways can be intervened. Our observations offer a unique first step to investigating the role of amino acid variability at mimicked functional sites that might help explain differences in phenotypic outcome associated with viral infections.

Finally, the repertoire of structural mimics we discover opens new opportunities to identify potential mechanisms underlying autoimmune disorders of viral origin and new protein-based immune-modulatory therapeutics. For example, leveraging mimicry relationships between coronaviruses and human proteins informs about cellular targets and signaling cascades that are tuned during infection. Such knowledge can provide important clues about pathways that mediate pathology associated with infection and may point the way for designer therapeutics directed at these pathways. In the short-term, and as highlighted by recent discovery of SARS-CoV-2 risk factors^31^, information about virus-encoded structural mimics can help refine large-scale clinical studies and reveal determinants of immunity, susceptibility, and clinical outcome associated with human infection.

## ACKNOWLEDGEMENTS

We thank Donald Petrey for technical support. This work was funded by NIH grants 5R01GM109018-05 and 5U54CA209997-03 to SS, 5R01GM030518-39 to BH, 5R01GM117591 to RR and equipment grants S10OD012351 and S10OD021764 to the Columbia University Department of Systems Biology.

## MATERIALS AND METHODS

### Viral dataset assembly

We compiled a dataset of 7,486 viruses, represented by a total of 337,493 viral protein amino acid sequences together with annotations of their corresponding hosts, from virus-hostDB as of October, 2016^35^. Viruses were classified according to the taxonomic divisions of their corresponding host based on NCBI taxonomic divisions^36^ (Table S1).

### Structural neighbor search

The 337,493 viral protein amino acid sequences were reduced to 269,077 non-redundant sequences filtered at 100% sequence identity with Cd-hit^37^. Sequence-based methods were then used to identify experimentally determined protein structures that, based on their sequence relationship to the viral protein, are expected to have similar structures (referred to here as “structural templates” - Figure 1a). First, viral proteins were parsed into domains using CD-search^38^. Next, to identify structural templates, full sequences and parsed domains were queried against the PDB^39^. Sequence homology search was performed in three steps, where each step is run only if the preceding step fails to report a structural template with a conservative E-value < 1×10^−12^: i) Blast-based search ^40^; ii) HHblits-based search^41^ and; iii) the third step runs five iterations of PSI-Blast. The pipeline reports the best structural template mapping to non-overlapping sequence segments for each query protein. When multiple structural templates map to the same segment in a query protein, only the structural template(s) with the lowest E-value are reported. The set of structural templates, describing the structural space of viral proteins, is further refined in order to minimize the number of redundant templates. To this end, structural templates derived from the same pdb chain and with their start and end positions within 15 residues are clustered together, using the longest structural template as the cluster representative. Finally, the set of structural templates is searched against the PDB database with Ska^12,13^ using a structural alignment score (SAS) < 2.5Å as a cut-off to identify structurally similar proteins^15^. The ska algorithm focuses on alignment of secondary structure elements and uses only C-alpha coordinates of residues. Hence any differences in loop or side-chain conformations that might be expected because of differences in resolution would have little to no effect on a ska alignment.

### Sequence homology assessment between viral proteins and their corresponding structural neighbors

Sequence homology among a pair of structurally similar proteins was considered significant when neither a Blast, HHBlits or PSI-Blast alignment yielded an E-value < 1×10^−6^. In addition, for figures 2 and 3, sequence identities for pairs of structurally similar proteins, were computed using the Needleman-Wunsch algorithm implemented in the EMBOSS package (considering only the corresponding protein segments sharing structural similarity)^42^.

### Calculating overlap between structural space of a virus and proteins in a given taxonomic division

Taxonomic division enrichment (Figures 1b-c) was computed with a hypergeometric test that describes the significance of having *k* structural neighbors belonging to a particular taxonomic division (out of *n* total structural neighbors for a group of viruses) given the entire set of structurally solved proteins in the PDB of size *N* that contains *K* proteins from the same taxonomic division^43^. To minimize experimental bias of multiple PDB entries for the same protein, structurally solved proteins were mapped to their Uniprot accession codes^44^ using SIFTS mapping^45^.

### Protein modeling and structural neighbor search for *Aedes aegypti* proteome

Our validated, inhouse modeling pipeline^1,8^ was used to model the 16,652 protein sequences (obtained from the Uniprot database^44^) in the *Aedes aegypti* proteome. Three-dimensional models for full-length proteins and protein domains, as defined by the Conserved Domain Database^38^, were either taken directly from the PDB^39^ or built by homology modeling as described previously^8,11^. Protein structure models were built using NEST^12,46^. Structural neighbor search was run on the set of modeled mosquito proteins using Ska^12,13^ and a structural alignment score (SAS) < 2.5Å was used as a cut-off to identify structurally similar proteins^15^. The modeling and structural neighbor search pipeline reported structural neighbors for 10,895 (65%) mosquito proteins.

### Identifying the mosquito structural neighbor space for each virus in the dataset

The unique set of mosquito structural neighbors was computed for across all viruses. Structural relationships between viral and mosquito proteins was inferred by identifying common structural neighbors shared between a given viral and mosquito protein pair. Poorly annotated viruses with less than 4 annotated viral proteins were discarded. Invertebrate-infecting viruses were further sub-classified into i) invertebrate-infecting viruses (including mosquito infecting viruses); ii) viruses infecting both human and mosquitoes and; iii) viruses infecting mosquitoes but not humans. We applied the non-parametric Mann-Whitney test to compare the size of the mosquito structural neighbor space per virus (normalized by the total number of annotated viral proteins in each virus) between different groups of viruses (Figure 1c), and outliers were removed with the Rout method (Q = 1%) implemented in Prism^47^.

### Human protein structural neighbors of human-infecting viral families

To estimate the number of human protein structural neighbors per virus, viruses with 4 or more annotated proteins with structural template for at least 30% of the viral proteins were considered (Figure 2a, 2c). The total number of human structural neighbors per virus was corrected by the corresponding structural coverage (e.g. for a virus where 70% of its proteins have a structural template and shows structural similarity to 30 human proteins, the corrected number of human structural neighbors will be 30 / 0.7 = 42.9, Figure 1a). Viruses were grouped into viral families and families with less than 5 viruses were not plotted. In order to plot the number of human structural neighbors per residue for a given family of human infecting viruses (Figure 2c), we normalized the total number of human mimics per virus (Figure 2a) by the corresponding proteome size (Figure 2b).

### Functional enrichment analysis

Enrichment of biological ontologies (molecular function, biological pathways and diseases) was determined using David^48,49^, background corrected using the 5,841 human protein structures extracted from PDB. A Bonferroni corrected p-value < 0.01 was used to identify enriched biological ontologies. Viral families and biological ontologies enriched in at least one viral family were clustered using R and the heatmap.2 function within gplots package (values in original matrix: -log_10_ P-value_Bonferroni_; distance metric: Euclidean, method: complete) ^50,51^.

## FIGURE LEGENDS

**Figure S1.**
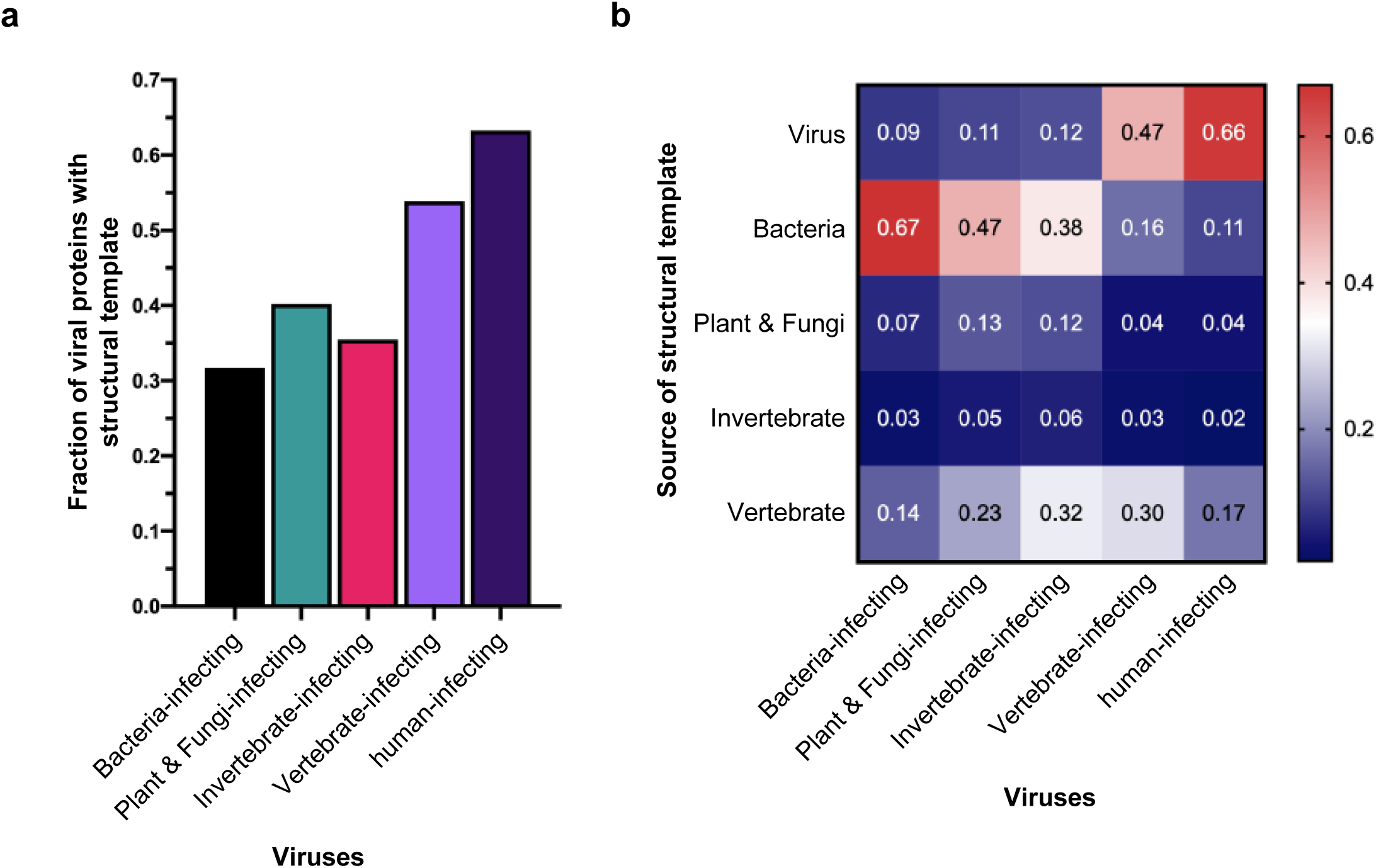
Structural coverage of viral proteins. **a**, Fraction of viral proteins, grouped based on the taxonomic division of the corresponding host, with a sequence homolog whose structure is experimentally known (structural template). **b**, Fraction of the structural templates belonging to a particular taxonomic division identified for different groups of viruses.

**Figure S2.**
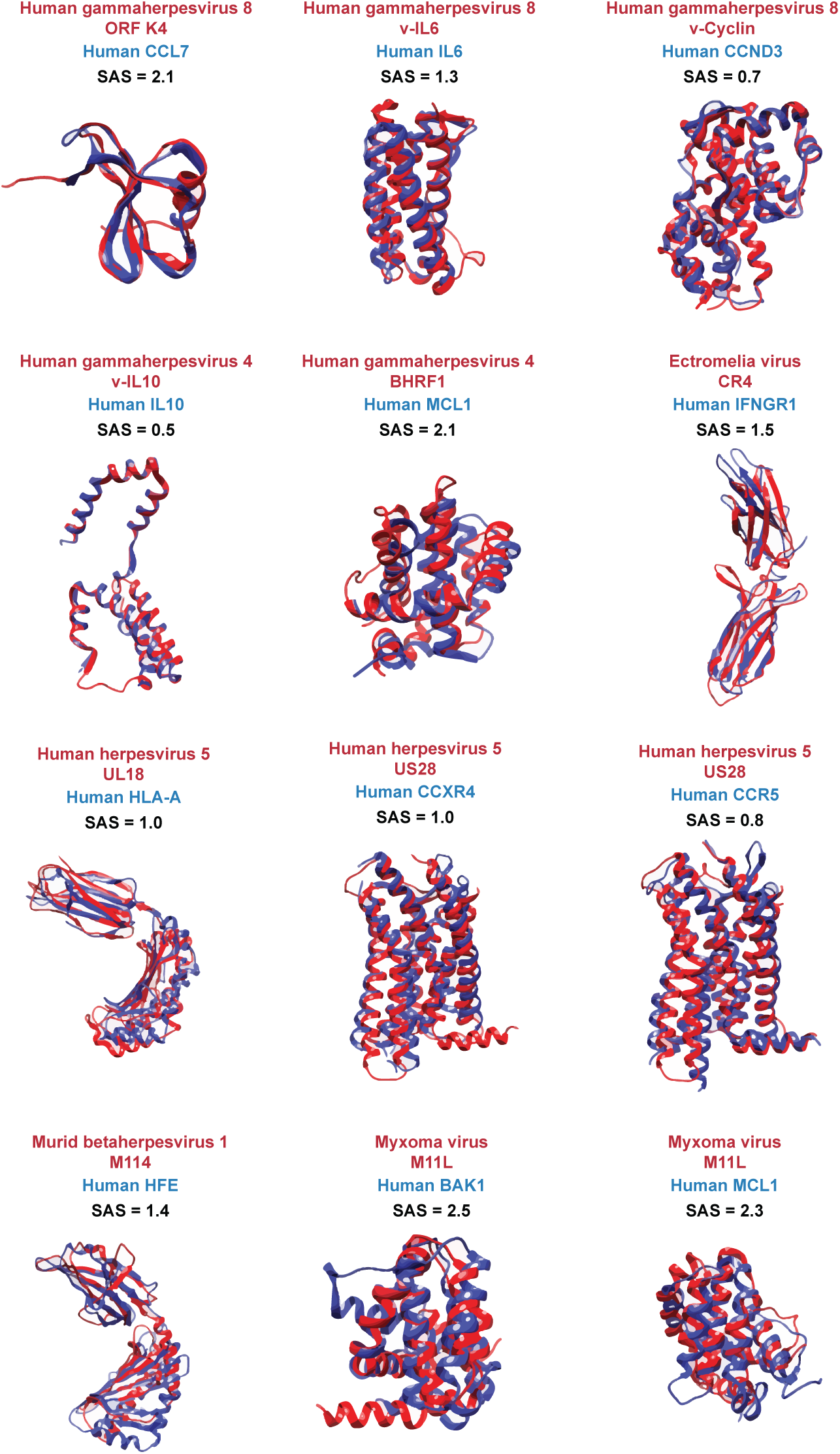
Virome-wide scan rediscovers known viral-host structural mimics. Shown are examples of known viral-host structural mimics^5,32,52-55^ recapitulated with structural neighbor search. Viral and host proteins are colored in red and blue respectively. SAS is shown in black.

